# COATswga: A Coverage Optimizing and Accurate Toolkit for fast primer design in selective whole genome amplification

**DOI:** 10.1101/2025.11.26.688640

**Authors:** Kaleb Zuckerman, Alec Leonetti, Rebecca DeFeo, Om Taropawala, Alfred Simkin, Abebe A. Fola, Jeffrey A. Bailey

**Affiliations:** Center for Computational Molecular Biology, Brown University, Providence, RI, USA; Department of Pathology and Laboratory Medicine, Brown University, Providence, RI, USA; School of Public Health, Brown University, Providence, RI, USA

**Keywords:** sWGA, COATswga, primer design, *Plasmodium falciparum*, pathogen enrichment, WGS

## Abstract

**Background:** Despite the transformative nature of next-generation sequencing in genomics, efficiently capturing underrepresented microbial DNA from complex biological mixtures, such as pathogens from host samples, remains a persistent challenge. Unwanted genomes make whole-genome sequencing (WGS) inefficient and expensive without enrichment of the targeted genome. Selective whole genome amplification (sWGA) uses primers that selectively bind to the target genome to preferentially amplify and enrich an entire microbial genome. However, existing sWGA primer design is often complicated and time-consuming, with primers usually having biased amplification producing uneven genome coverage. Developing faster primer design methods that ensure uniform, reliable microbial genome recovery are critical to improving sWGA.

**Methods:** We developed COATswga, a Coverage Optimizing and Accurate Toolkit for designing sWGA primer sets. The pipeline consists of three key steps: (1) *primer discovery*, using k-mer counting to identify all candidate primers in the target genomes; (2) *filtering*, where primers are screened for amplification potential, thermodynamic stability, and specificity; and (3) *set formation*, which uses a novel interval-based tiling algorithm. COATswga is parallelized for efficiency and supports pre-existing primer set refinement and gap filling. The pipeline was evaluated by designing primer sets for *Plasmodium falciparum* (strain 3D7) and benchmarking their performance against sets generated using similar pipelines.

**Results:** COATswga-designed primers demonstrated superior enrichment based on both qPCR and long-read sequencing. In *P. falciparum*-spiked human DNA samples at parasitemia levels of 400, 100, and 10 parasites/μL, COATswga primers achieved higher amplification efficiency and specificity compared to alternatives. At 100 parasitemia, COATswga yielded ∼99% of reads mapping to the *P. falciparum* genome with 82.5% of the genome having 5x coverage. Even at 10 parasites/uL, COATswga primers enabled effective amplification of key drug resistance genes (*pfcrt* and *pfmdr1*). Additionally, integration with molecular inversion probe (MIP) genotyping showed that COATswga greatly improved sequencing depth and target recovery in low-density infections compared to established sWGA primers.

**Conclusion:** COATswga provides a robust and flexible solution for designing sWGA primers to reliably recover targeted genomes from complex samples. The new *P. falciparum* primer set performs well in low-parasitemia samples allowing more extensive application of sWGA in falciparum malaria genomics.

## Background

In the last few decades, advancements in next-generation sequencing (NGS) technologies have revolutionized science, enabling large-scale affordable genomic studies [1]. Despite this, the study of specific microbes in the variable milieu from which they originate remains challenging [2]. Their small genomes that should make microbes inexpensive to sequence are often found in complex mixtures of other microbes and/or larger amounts of host genome [3]. All of this makes targeted microbial DNA sequencing inefficient and costly. Moreover, with less than 1% of known microbes successfully cultured, the need for tools to specifically capture whole genomes from variable assortments of organisms becomes even more crucial [2]. To address these challenges, several methods have been developed to isolate target microbial DNA and improve whole genome sequencing (WGS), including hybrid capture, single-cell analysis, and selective lysis [4–6], though these approaches can be infeasible due to the higher cost and assay complexity.

Selective whole genome amplification (sWGA) of a target pathogen offers a promising solution. This cost-effective and single-tube technique enables the amplification of entire target genomes from mixtures, avoiding extensive purification or culturing [7]. By designing sets of DNA primers that bind selectively to the target genome (the foreground) while minimizing amplification of any contaminating DNA (the background), sWGA can exponentially increase the amount of target DNA until it predominates a sample. Due to the large number of potential primers in even a small microbial genome, sWGA primer sets are selected by software that identifies DNA motifs occurring frequently in the foreground (targeted genome) and rarely in the background (‘contaminating’ genome(s)). Amplification relies on Phi29 DNA polymerase, which offers high fidelity branching amplification, with an error rate 100 times less than Taq, and high processivity, producing up to 70-kbp fragments ideal for studies of whole genomes [8, 9]. Recent innovations, such as the EquiPhi29 polymerase, have further improved this process by enabling the use of longer and more complex primers at elevated temperatures [10]. sWGA has already been successfully applied to various pathogens, including *Mycobacterium tuberculosis*, *Wolbachia pipientis*, *Plasmodium falciparum, Plasmodium vivax*, *Neisseria meningitidis*, *Coxiella burnetii*, and *Wuchereria bancrofti*, among others [3, 11–14].

Despite sWGA’s successes, computational tools for rapidly designing effective primer sets for sWGA have lagged. Effective primer design requires maximizing specificity while still evenly tiling the target genome to avoid over-amplifying specific regions or leaving gaps in coverage. Existing tools, such as the swga toolkit [3] and swga2.0 [15], have made significant progress in this area but still exhibit limitations in runtime efficiency, uniformity of target genome coverage, and adaptability to different experimental conditions such as variable length primers.

To further improve sWGA primer design, we developed the COATswga pipeline. COATswga leverages an interval-based algorithm and a highly parallelized structure to provide primers that improve genome coverage and selectivity, all while drastically reducing runtime compared to other similar pipelines. We evaluated COATswga by designing a primer set for *Plasmodium falciparum*, experimentally comparing their performance with the primers generated by swga2.0 augmented with hand-selected primers, and the currently used primer set for selectively amplifying *P. falciparum*, the Sanger set [16]. *P. falciparum* is the most prevalent species responsible for over 90% of malaria cases and deaths in sub-Saharan Africa and is often at low abundance in samples, with human DNA representing the vast majority of DNA isolated from blood. Our results, assessed using qPCR and Oxford Nanopore sequencing, demonstrate that the COATswga primer set offers superior selectivity, coverage, and practical utility, positioning it as a valuable tool for sWGA primer design.

## Materials and Methods

### The COATswga Pipeline

The structure of the COATswga pipeline is divided into three main steps **(Figure 1)**. The program searches the foreground and background genomes provided for all possible primers, filters out ineffective candidates, and forms sets of the most optimal primers. The top sets found are outputted to the user.

**Figure 1:**
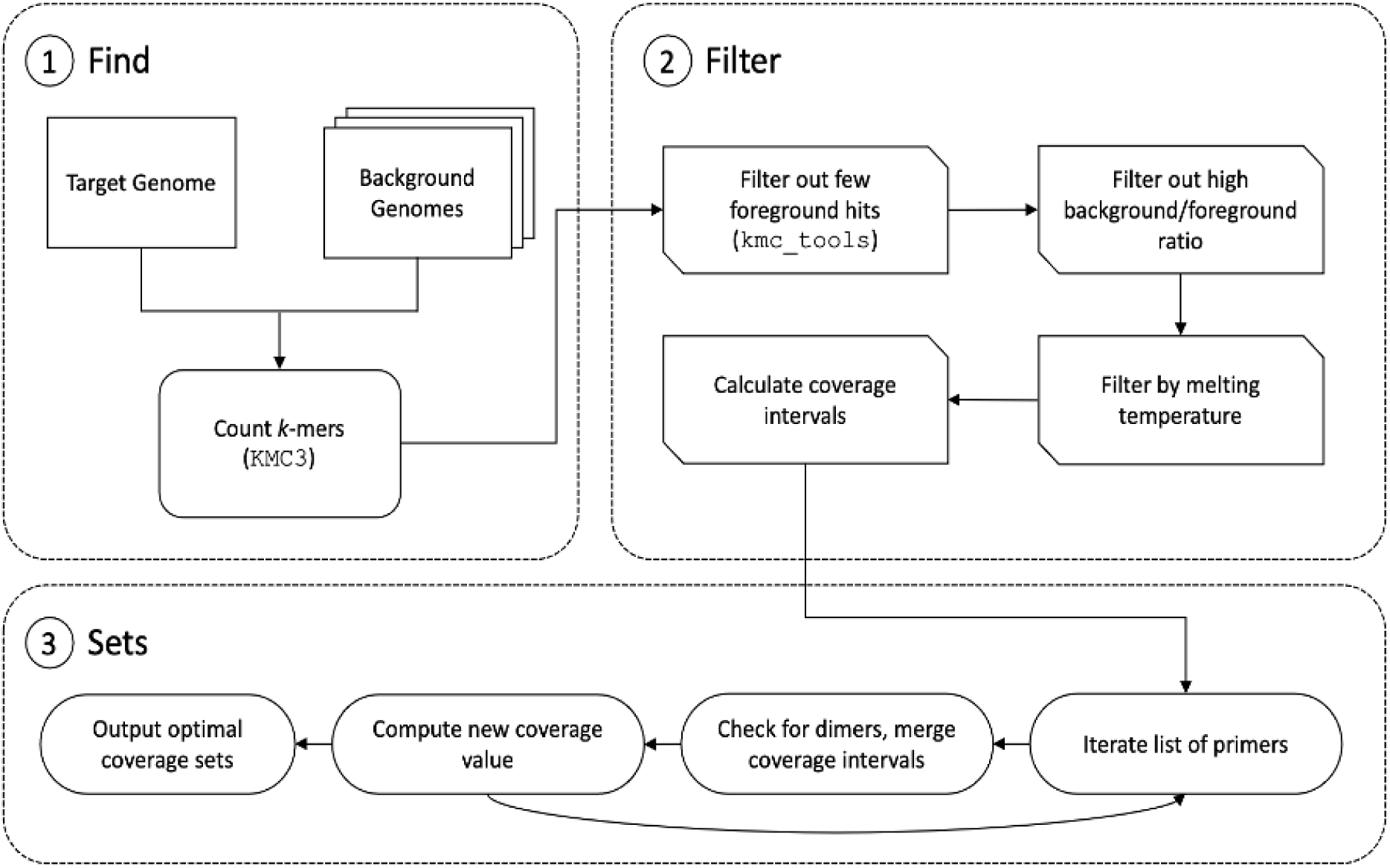
An overview of the COATswga workflow. The pipeline works in three steps. It begins by counting all k-mers with KMC3, then filtering those motifs that do not meet the criteria specified by the user (foreground frequency, background-to-foreground ratio, melting temperature, etc.), finally forming sets of primers that do not form self- or primer-primer dimers and sufficiently tile the target genome.

#### Step 1: Find

The pipeline starts by finding all possible primer candidates. All *k*-mers in the foreground and background genomes are identified and counted for the parameters min_primer_length ≤ k < max_primer_length, using the disk-based *k*-mer counter KMC3 [17]. Here, COATswga can handle multiple foreground and background files, allowing for the processing of complex DNA profiles. K-mer counts are saved in the specified directory (data_dir) using the given foreground and background prefixes, only running KMC3 if the files do not already exist. This allows for rapid repeat runs of the program for fine tuning of the other selection parameters, as the k-mer counting step is often the most time-consuming.

#### Step 2: Filter

After the set of all candidate k-mers has been identified, potential primers that would be ineffective must be filtered out. COATswga leverages kmc_tools to quickly remove primers that bind too infrequently to the foreground (<min_fg_count), thus having low amplification potential, or too frequently to the background when compared to the foreground (>max_ratio), thus having poor selectivity. It then removes primers with melting temperatures outside the specified range (>min_tm, <max_tm), as calculated by melt (https://pypi.org/project/melt/). The primers are then sorted by a combination of the ratio of the number of background hits to the number of foreground hits (referred to as the background-to-foreground ratio or simply the ratio) and the foreground frequency, so the most effective potential primers are considered first. This step is parallelized at multiple points to ensure maximum efficiency. Importantly, the primer set will be more specific the lower the ratio is, with a ratio of zero indicating perfect specificity to the target.

#### Step 3: Sets

Once the suboptimal primers have been removed, COATswga transitions to collecting position information to prime the set-forming algorithm. It parses the foreground genome, recording the positions of each primer location on the forward and backward strands. It then passes those positions into bedtools [18], making interval calculations to precompute the coverage intervals of each potential primer and storing those intervals in a dictionary for fast access.

To form sets, the algorithm iterates through the list of potential primers, checking for potential self- or primer-primer dimers using an algorithm adapted from PrimerROC [19], as implemented by Jason Hendry [20]. If the primer passes the check, the coverage interval of the primer is merged with the coverage intervals of the current set, calculating the change in genome coverage. A threshold of the percent of new bases covered must be passed for the primer to be added to the set. The threshold is stepped down with each iteration, requiring the highest amount of novel genome coverage in the first iteration of the primer list, decreasing with each subsequent iteration. This ensures that the most impactful primers are added first before filling in coverage gaps.

The algorithm continues cycling through the primer list until it either fully covers the minimum proportion of the genome specified by the user (target_coverage) or runs out of primers to check at the lowest threshold. In this way, the algorithm ensures proper tiling of the target genome, guaranteeing even coverage. It runs the set-forming algorithm multiple times in parallel, with each run initializing the set with a different primer. Importantly, the set forming algorithm is run first on the forward strand and then the reverse strand to ensure exponential amplification. Of all sets formed, the sets with the lowest ratio, highest predicted coverage, and fewest number of primers in the set are outputted to the user, along with the predicted forward and reverse coverage, the number of hits in the foreground and background genomes, and the ratio for each set. There is a verbose option to print out all sets considered.

### Methodology for the coverage calculation

The algorithm for determining primer set membership efficiently replicates the mechanism of DNA amplification to give a more accurate estimate of coverage. In DNA replication, the polymerase enzyme starts at each primer and synthesizes DNA for a number of bases, thereby covering that section of the genome. In the same way, the algorithm takes the positions of each primer and considers the section of the genome after the primer position as “covered” for the fragment length specified by the user (meant to represent the average fragment length of the enzyme being used). It stores this segment as an interval with a start and end position, reducing memory usage and enabling the use of bedtools for rapid interval arithmetic.

In addition to generating primer sets *de novo*, COATswga has a gap-filling functionality, enabling it to complement primer sets currently in use for sWGA. Within the parameters of the pipeline, there is an option to specify primers already in an existing set (existing_primers). During the set-forming step, the algorithm ensures that any primers added to create the final sets will not form primer-primer dimers with those in the existing set. In this way primer sets that already perform well but miss a few key areas can be improved to better cover gaps or regions of low coverage without attempting an entirely new design. t

### Experimental Testing of sWGA Primer Sets

#### Protocol for the sWGA reaction

To test the specificity and efficiency of the designed primers, DNA samples were prepared by spiking *P. falciparum* DNA into extracted human DNA to simulate infected blood samples at approximate parasitemias of 400, 100 and 10 parasites/uL. The DNA samples were then subjected to a modified selective whole genome amplification (sWGA) protocol. First, we combined 8 µL of DNA sample, 0.25 µL (20 µM concentration) of the primer pool, 0.5 µL of 10X ThermoFisher EquiPhi29 reaction buffer, and 1.25 µL of nuclease-free water, bringing the total volume to 10 µL. The mixture was denatured at 95°C for 3 minutes. Next, we added 1 µL (10 units) of EquiPhi29 DNA polymerase, 2 µL of reaction buffer, 0.2 µL of 100 µM DTT (used to stabilize the enzymes), 2 µL of 10 mM dNTPs, and nuclease-free water to reach a total pool volume of 20 µL (DL-Dithiothreitol; Clelands reagent). This mixture was then incubated at 45°C for 3 hours (isothermal amplification), followed by 65°C for 10 minutes to deactivate the enzymes.

### Qubit (total DNA) and duplex qPCR (targeting *P. falciparum* and human)

Amplification was confirmed with a Qubit Fluorometer (Thermo Fisher) according to the manufacturer’s protocol. Specificity of amplification was initially assessed with duplex qPCR analysis of the amplified samples targeting human and *P. falciparum*. For the qPCR, 2 µL of amplified DNA sample was mixed with 1.22 µL of molecular-grade water, 6 µL of iQ Supermix (Bio-Rad laboratories), 0.06 µL of 10µM of the HumTuBB F and R primers and probe, 0.24 µL of pfldh F and R primers, and 0.12 µL of pfldh probe. Samples were plated on Bio-Rad qPCR plates and run on a Bio-Rad CFX96 Real-Time System with C1000 Thermal Cycler base. Samples that amplified effectively were then sequenced with Oxford Nanopore Technologies for more in-depth analysis.

### Whole genome sequencing using Oxford Nanopore Technologies (P2)

Amplified products were debranched using NEBuffer 2 and T7 Endonuclease I (New England BioLabs) and cleaned with Sera-Mag beads. The samples were then sequenced on Oxford Nanopore platforms (P2). Basecalling was performed by the Oxford Nanopore Guppy basecaller algorithm (now deprecated, Oxford Nanopore recommends using Dorado, found at https://github.com/nanoporetech/dorado). Reads were mapped to their respective genomes with Minimap2 [21] and indexed with samtools [22]. We used qualimap [23], IGV [24], bamcov [25], and the R programming language [26] for additional data analysis and visualization.

### MIP sequencing and data analysis

We tested newly designed COATswga primers to enrich targeted DNA before molecular inversion probe (MIP) genotyping. sWGA amplified DNA samples (as described above) were targeted, captured, and sequenced using MIPs designed to cover key *P. falciparum* resistance genes associated with artemisinin and partner drug resistance, including *pfkelch13*, *pfmdr1*, *pfcrt*, *pfdhfr* and *pfdhps* genes. MIP capture and library preparation were performed as previously described [27, 28] with some modification. The sWGA amplified product was subjected to 4x dilution and 5ul of diluted product was used as a template for MIP capturing. MIP sequencing was conducted using an Illumina NextSeq 550 instrument (150 bp paired end reads) at Brown University (RI, USA).

The raw data generated using MIPs were demultiplexed using MIPtools software (https://github.com/bailey-lab/MIPTools), which is a computationally suitable tool for MIP data processing and analysis. The data was further processed using MIP Wrangler software (https://github.com/bailey-lab/MIPWrangler), in which sequence reads sharing the same Unique Molecular Identifiers (UMIs) were collapsed to generate a single consensus. MIPwrangler calls micro haplotypes whose abundance per sample was assessed for each MIP relative to other samples and across MIPs. Each dataset was analyzed by mapping sequence reads to the *P. falciparum* 3D7 reference genome using Burrows-Wheeler Aligner (BWA) to generate a total number of sequenced reads per sample and sequencing coverage per sample per MIP probe used. Comparison of coverage and read depth across different parasite density were determined for each sample set (before and after sWGA) using R software and *p*-value of ≤ 0.05 was considered statistically significant.

## Results

### Primer design for P. falciparum with COATswga

We used COATswga to design a primer set for *Plasmodium falciparum* (strain 3D7, GCF_000002765.5) based on a background of *Homo sapiens* (GRCh38.p14). We ran the COATswga pipeline with a target coverage of 97%, a temperature range of 15°-45°C to reflect our use of the improved EquiPhi29, and a fragment length of 10 kilobases, which is a conservative view of the performance of Phi29. Between multiple runs, we increased the minimum foreground count and the maximum background-to-foreground ratio to find the most selective set that would still give even amplification. We chose a set of 13 primers (**Table S1**) with a ratio of 0.52 and an average expected coverage of 96%. We verified these numbers by searching the genome with both grep and a custom search algorithm to find all matching motifs. Even coverage was confirmed by visualizing with IGV (**Figure S1**).

### Primer design of the swga2.0 set

For a direct comparison, we also used the swga2.0 pipeline to generate a primer set for the same foreground (*P. falciparum*) and background (human). The swga2.0 produced 10 primer sets with 3-6 primers in each. Concerned about under-amplification due to the small number of primers in each set from swga2.0 and given that almost all the sets reused some of the same primers, we formed a superset of the primers from all 10 sets (dropping two that formed primer-primer dimers) in addition to some hand-picked primers with low ratios and Gini indexes, for a total pool of 25 primers (**Table S1**).

In addition to the swga2.0 pool we used the currently accepted sWGA primer set containing 10 primers for amplifying *P. falciparum*, referred to as the Sanger set [16], to amplify the same DNA samples as a benchmark for both newly designed primer sets.

### Comparison of qPCR results between primer sets

The qPCR results of the sWGA products with *Pfldh* primers confirmed successful amplification of *P. falciparum* from all three sWGA primer sets, also indicating that the COATswga primers gave the best specificity (highest *P. falciparum* to human ratio) for all three of the mock parasitemias (**Figure S2**). The average *P. falciparum* fold change (ratio of copy pre and post sWGA ) was greatest with the COATswga primers in the 100 parasitemia samples, followed by the augmented swga2.0 pool. COATswga and the Sanger set performed similarly for the 400 parasitemia samples. Though the yield for the Sanger set was greater at the 10 parasitemia level, it also amplified the human DNA over 10,000 times more than the COATswga pool. The swga2.0 pool gave the lowest fold change for both the 400 and the 10 parasites/uL (**Figure S2**).

### Comparison of genomic coverage between primer sets with NGS

Ultimately, sequencing is needed to fully assess sWGA performance. We sequenced the 100 parasite/uL samples for each primer pool as well as the COATswga-amplified 10 parasites/uL sample, to test performance on low parasitemia samples. swga2.0 did not show enough enrichment to warrant sequencing at 10 parasites/uL level, so we did not sequence the other two pools for comparison. For analysis at the 100 parasitemia level, the number of sequenced bases used for analysis was normalized to 1.1 gigabases (Gb) per sample (including both human and *P. falciparum* DNA), the lowest amount of the three. Analysis of the COATswga 10 parasitemia sample used 538 megabases (Mb) of sequencing data.

Oxford Nanopore sequencing of the sWGA products at 100 parasite/uL for each of the three primer sets confirmed the qPCR results, revealing that the COATswga primers were the most selective and gave the best coverage profile. COATswga had 98.9% of reads mapping to the *P. falciparum* genome with a mean coverage of 49.9 and 82.5% 5x coverage. The Sanger set had a 95.4% mapping ratio, with a mean coverage of 45.2 and 78.3% 5x coverage, and the swga2.0 pool had 55.5% of reads mapped to *P. falciparum*, a mean coverage of 16.3 and 78.5% 5x coverage (**Figure 2A**). COATswga also gave the highest coverage at each depth >3x by several percent, although the augmented swga2.0 pool gave the highest 1x coverage, covering 2% more of the genome compared to COATswga (96.2% vs 94.2%) (**Figure 2B**). At the 10 parasitemia level, sequencing data showed COATswga gave 60% 1X coverage and 32.5% 5X coverage with 94.5% of reads mapping to *P. falciparum*. Importantly, it gave sufficient coverage for study of two key drug resistance genes in *Plasmodium falciparum*, *pfcrt* and *pfmdr1* (mean coverage 4x and 10x, respectively).

**Figure 2:**
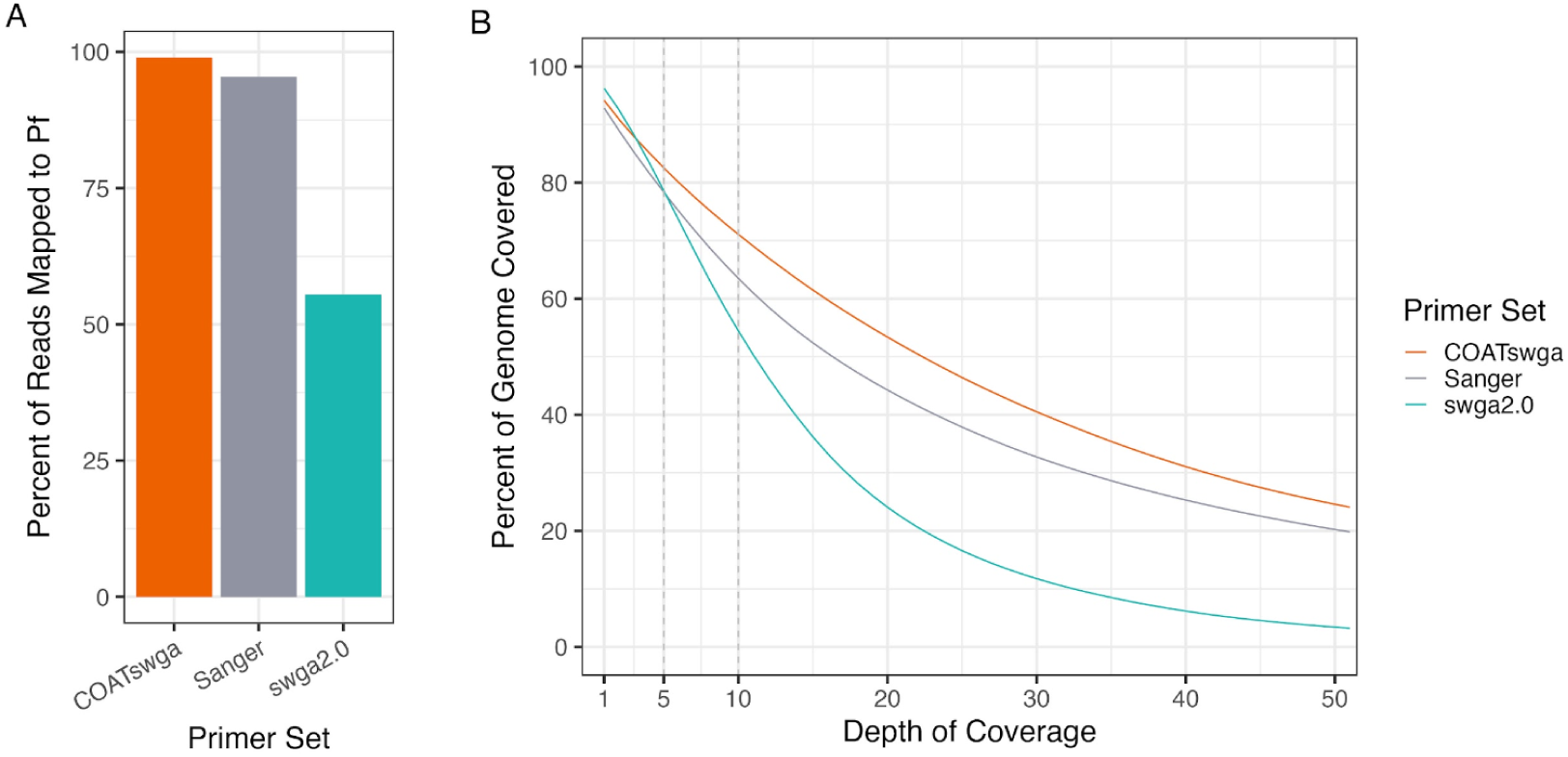
Deep sequencing (1.1 Gb of sequencing data for each primer set) of the 100 parasitemia sWGA experiments with the four primer sets. (A) Bar graph of the selectivity of the three primer sets from the sequencing data reveals that the COATswga primers were the most selective. The y-axis represents the percent of reads mapping to the *Plasmodium falciparum* genome, the x-axis represents the primer set. (B) Plot of the percent of the genome covered at each depth from 1x to 50x shows how the COATswga primers give the best coverage at higher depths.

To assess performance in which the costs of sequencing may be limiting, we analyzed genome coverage as a function of sequencing effort (**Figure 3**). Results showed that over 350 Mb of data were required for the swga2.0 pool to surpass COATswga in 1x coverage. Below this threshold, COATswga consistently outperformed the other sets in 1x, 5x, and 10x coverage. The Sanger set also outperformed swga2.0 at higher coverage depths and maintained superior 1x coverage up to 250 Mb of sequencing data.

**Figure 3:**
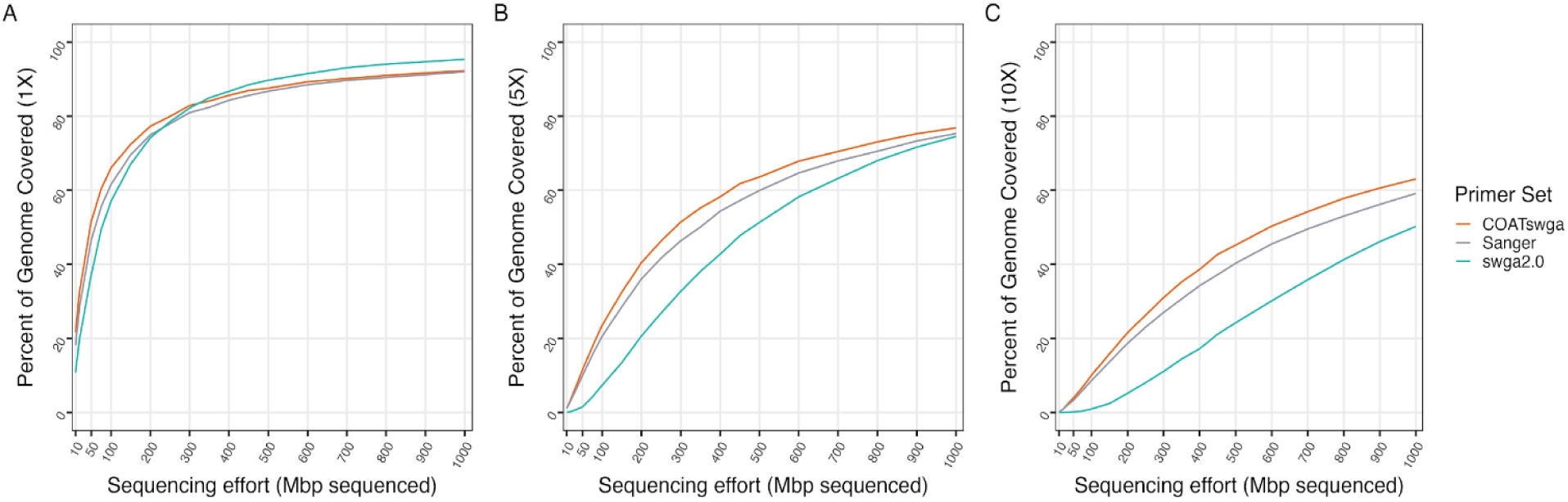
Percent coverage of the genome per megabase of DNA sequenced for (A) 1X coverage (B) 5X coverage and (C) 10X coverage. (A) shows that relatively deep sequencing is required to get higher coverage with the `swga2.0` primers while `COATswga` outperforms handily in all other metrics.

A spatial coverage plot of the sequencing data (**Figure 4**) and bioinformatic analysis showed that all three primer sets generated similar areas of high and low coverage. Further investigation into whether combining different primer sets could enhance the depth or breadth of genomic coverage revealed that sequencing the same sample with multiple primer sets did not significantly improve coverage when compared to performing deeper sequencing with just the COATswga primers (**Figure S3**). This indicates that all three primer sets fail in similar areas, leaving room for further optimization.

**Figure 4:**
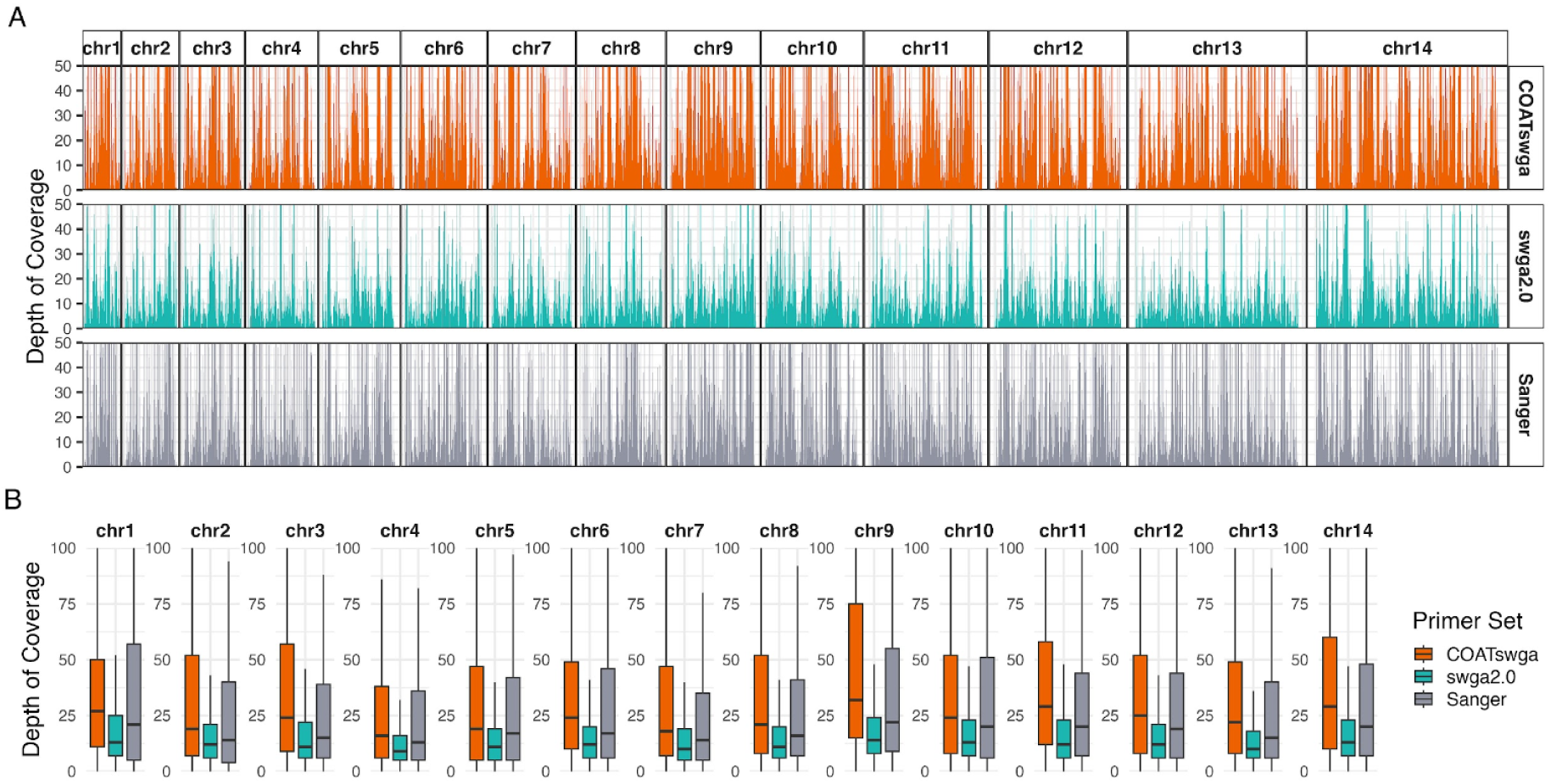
Depth of coverage across *P. falciparum* genome. (**A**) Depth of coverage by position on the genome. All three primer pools have similar peaks and troughs in coverage throughout the *Plasmodium falciparum* genome. We hope to use the hole-filling functionality in the COATswga pipeline to fill in the low-coverage areas. (**B**) Box plot of

### Drug resistance and *var* genes coverage and depth analysis

Malaria surveillance, particularly in the context of drug resistance, often focuses on five genes, *pfdhfr-ts*, *pfmdr1*, *pfcrt*, *pfdhps* and *pfkelch13*, with *pfkelch13* mutations associated with partial resistance to artemisinin (ART-R) [29, 30]. A comparison between the three primer sets showed that COATswga primers outperformed the alternatives, offering superior coverage of both the drug resistance genes and the genome as a whole (**Figure 5**). To further validate these findings, we extracted key drug resistance loci from the five major antimalarial resistance genes (**Table S3**) and assessed their coverage. This analysis confirmed that the COATswga primers provide superior coverage of the drug resistance genes compared to the other primer sets (**Figure 5)**. Bioinformatic combinations of the primer sets showed again that deeper sequencing of the COATswga samples gave better coverage of the drug resistance genes compared with combining different primer sets (**Figure S4**). Interestingly, the augmented swga2.0 primer set demonstrated enhanced coverage of the 61 *var* genes that we identified (**Table S2, Figure S6**), which code for the *P. falciparum* erythrocyte membrane protein 1 (*PfEMP1*) key to blood vessel wall adhesion and are distributed across the *P. falciparum* genome [31, 32].

**Figure 5.**
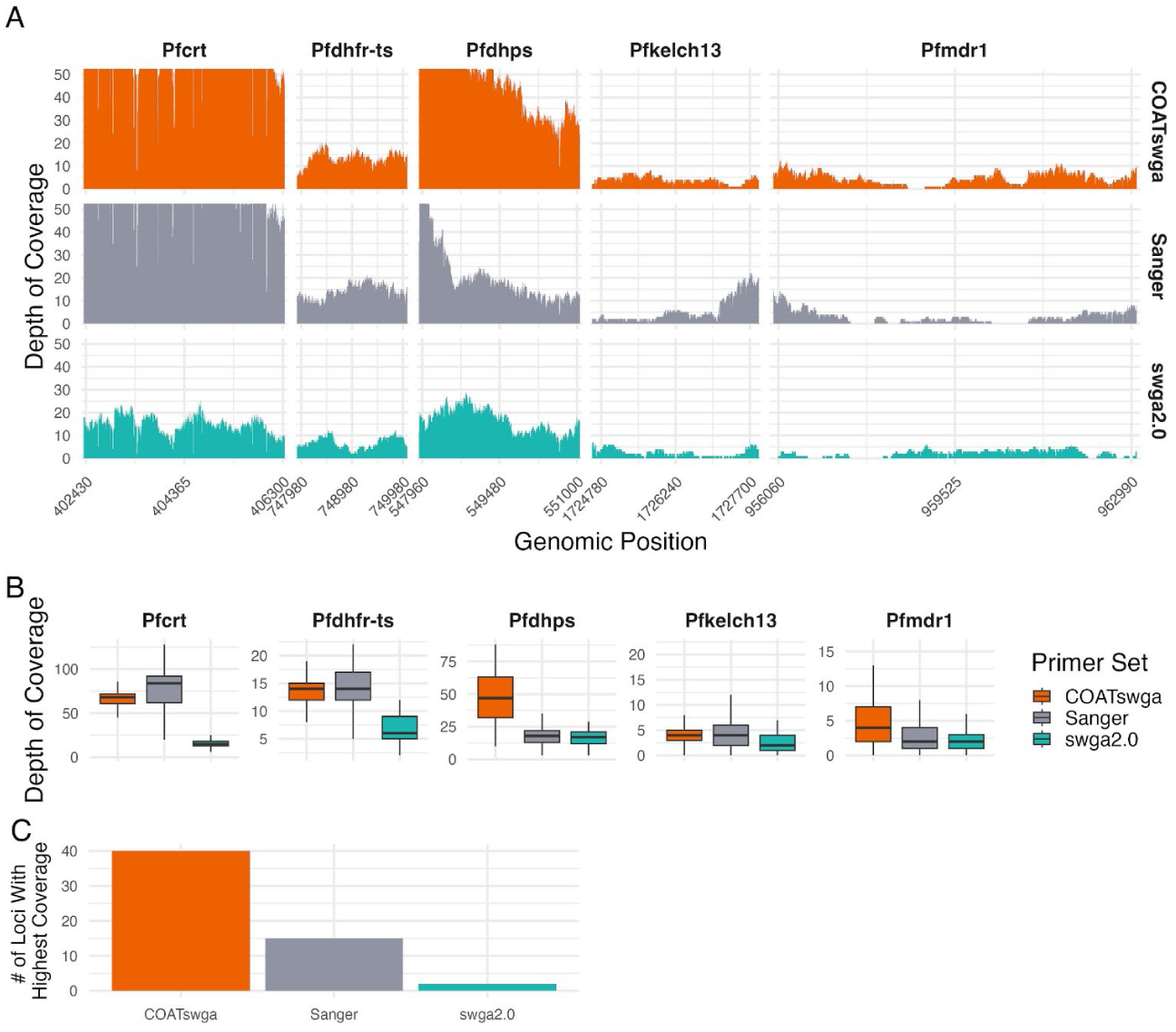
Depth coverage across key drug resistance genes. (**A**) Depth of coverage by position in each of the five key drug resistance genes. (**B**) Box plot of coverage values for the five drug resistance genes. COATswga

### Enhanced MIP sequencing through COATswga primers

We further evaluated the performance of our newly developed COATswga primers across a range of parasitemia levels as an input for targeted sequencing using Molecular Inversion Probe (MIP) sequencing. We observed a significant improvement in MIP sequencing performance when parasite DNA was first amplified using our custom COATswga primer set for selective whole genome amplification (sWGA). Compared to direct MIP sequencing without sWGA (**Figure 6A**), the inclusion of COATswga (**Figure 6B**) substantially increased sensitivity and coverage, particularly in low-parasitemia samples where DNA quantity is often limiting.

**Figure 6.**
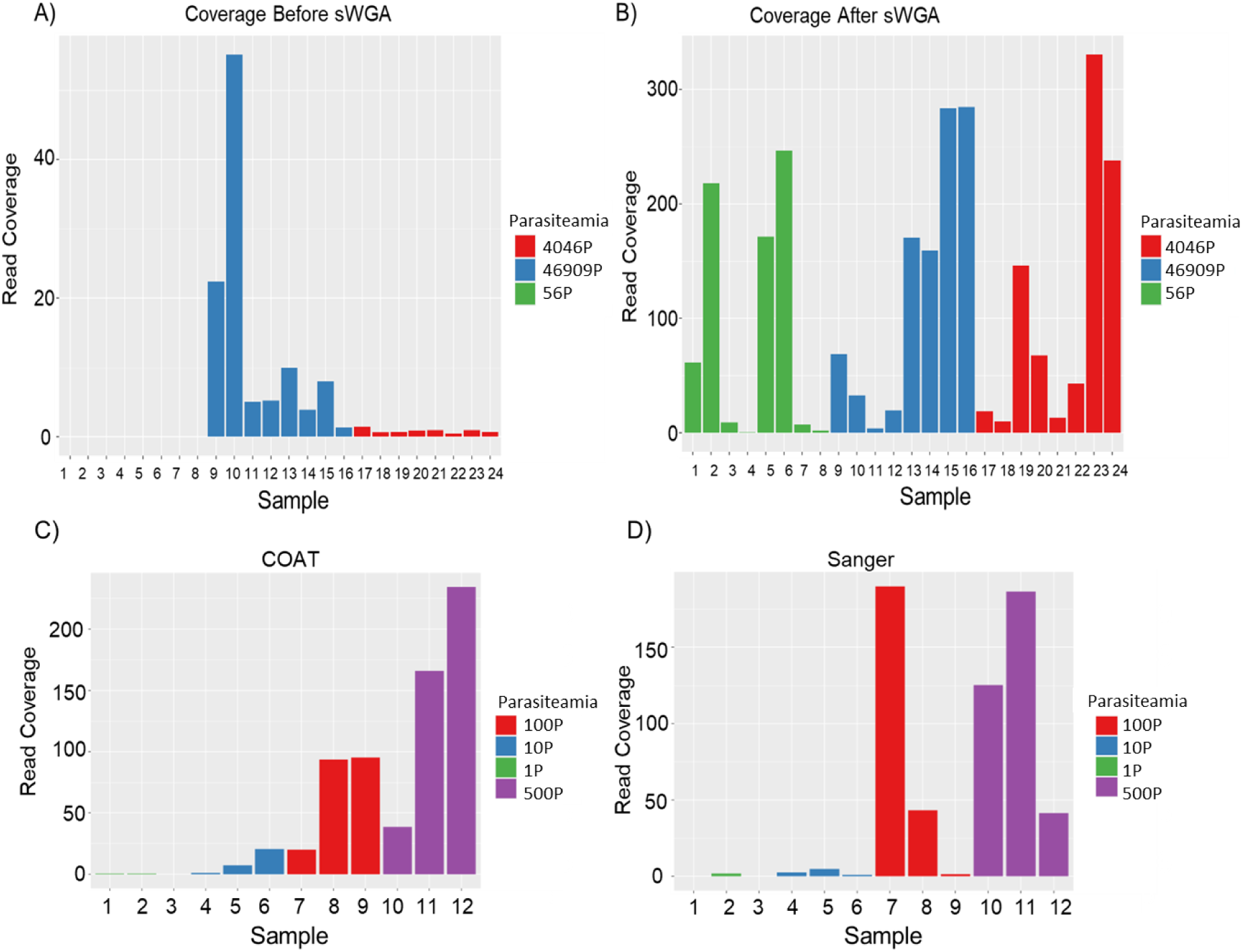
Evaluation of MIP Sequencing Performance. (A) Coverage obtained from MIP sequencing without COATswga pre-amplification. (B) Coverage following COATswga-based sWGA, demonstrating improved performance. Comparative analysis of MIP read coverage using COATswga primers (C) versus traditional Sanger primers (D), highlighting superior coverage with COATswga. P= parasites/uL

COATswga-based pre-amplification (**Figure 6C**) enabled more uniform and deeper MIP coverage, ensuring that even low-abundance parasite genomes could be reliably genotyped. This approach improved over non-amplified samples and those amplified with the widely-used current Sanger primers (**Figure 6D**).

Importantly, COATswga allowed successful genotyping at parasite densities as low as one parasite per microliter, a threshold at which conventional MIP sequencing or standard sWGA (sanger) methods typically fail. These improvements position COATswga-enhanced MIP sequencing as a powerful tool for molecular surveillance, especially in community surveys and elimination settings where asymptomatic low-density infections are common.

## Discussion

Our study presents a more flexible and rapid pipeline for sWGA primer design and an improved primer set for *Plasmodium falciparum.* For *P. falciparum*, compared to the existing Sanger [16] primer set and the swga2.0 [15] generated set that we augmented with additional primers, the COATswga primer set consistently demonstrated superior enrichment, and even coverage across the genome, especially in low-parasitemia samples. Oxford Nanopore sequencing at multiple parasite densities confirmed that COATswga achieved higher fold amplification of parasite DNA with minimal off-target amplification of human DNA. At the 100 parasites/μL level, COATswga showed exquisite specificity yielding nearly 99% of reads mapping to *P. falciparum* and the highest depth of coverage across multiple thresholds (>3x), while also delivering sufficient amplification down to 10 parasites/μL of key drug resistance genes such as *pfcrt* and *pfmdr1*. Base for base, it had better gene sequence coverage, outperforming both swga2.0 and the widely-used Sanger set. Importantly, coupling COATswga pre-amplification with Molecular Inversion Probe (MIP) sequencing significantly improved MIP SNP recovery and target depth, particularly at lower parasite densities where MIPs normally start to fail [33]. Thus our new *P. falciparum* COATswga primer set provides improved utility for malaria genomics and surveillance.

In this study, we highlight the critical factors that must be considered when designing effective primer sets for selective whole genome amplification (sWGA). Even for relatively small genomes like *Plasmodium falciparum*, millions of candidate primers can be computationally generated, creating an extensive search space. Effective narrowing begins by filtering on molecular attributes such as melting temperature (Tm), avoidance of secondary structures, and specificity. Primers that bind weakly to the parasite genome or excessively to human DNA, ubiquitous in clinical samples, must be discarded [16]. Previous sWGA pipelines, including the original swga [3] and its successor swga2.0 [15], implemented scoring metrics such as the Gini index to assess primer binding distribution. While the Gini index helps measure individual primer uniformity, it cannot guarantee that a full primer set provides balanced genomic coverage, leading to inconsistent amplification in some cases. To address this, COATswga introduces a novel interval-based algorithm that mimics biological DNA replication: instead of evaluating primers individually, it assesses sets of primers based on their collective ability to tile the target genome. This strategy significantly improves prediction of amplification success by avoiding uneven coverage and genomic dropouts. COATswga also enhances performance through parallelized computation, improved masking of repetitive regions, and background-sensitive modeling, optimizing primer selection for complex samples like dried blood spots. These improvements are crucial for enriching low-density microbial DNA in field-collected samples, where maximizing target yield and minimizing off-target amplification are critical for downstream sequencing and genotyping.

Building on these algorithmic innovations, our results further highlight key empirical factors that influence amplification efficiency and coverage. Notably, COATswga is the only currently available pipeline that supports the generation of variable-length primer sets, which likely contributes to its superior performance. Furthermore, we determined that primer sets must contain at least 10 primers to achieve consistent amplification success; Sets with fewer primers frequently failed to amplify adequately. Importantly, COATswga includes a gap-filling function that iteratively refines primer sets by targeting underrepresented genomic regions. This feature is particularly valuable for improving coverage in challenging regions of genomes, such as those associated with antimalarial drug resistance and the highly polymorphic *var* gene family in *P. falciparum*. Collectively, these capabilities make COATswga a powerful and flexible tool for both high-resolution population genomics and targeted sequencing applications in malaria surveillance.

In addition to enhancing whole-genome sequencing, COATswga significantly improved downstream targeted sequencing using molecular inversion probes (MIPs). By increasing genome coverage in both high- and low-parasitemia samples, COATswga enabled reliable MIP-based genotyping even in cases where traditional sequencing approaches typically fail due to insufficient template DNA. Its ability to effectively amplify target regions prior to MIP capture ensured sensitive and accurate detection of parasite variants, including low-frequency drug resistance mutations. This improvement is particularly valuable for field-collected samples with variable parasitemia, where reliable genotyping is often compromised.

While the COATswga-designed primers outperformed existing sWGA primer sets in terms of sensitivity and specificity, it is not without limitations. The computational cost associated with iterative primer refinement and gap-filling remains substantial, particularly for primer sets with fewer than 10 candidates, where multiple rounds of optimization may be required. Additionally, although the algorithm is designed to automate key aspects of primer design, fine-tuning often still requires manual parameter adjustments, which may increase runtime and reduce scalability in large population-based studies. Despite these challenges, the increased amplification performance and adaptability of COATswga make it a promising and robust tool for parasite genomic surveillance in diverse epidemiological settings.

Based on our experience, we recommend several strategies for optimizing primer design with COATswga. Starting with a more stringent maximum background-to-foreground binding ratio helps ensure primer specificity, with gradual relaxation if primer sets are insufficient. The pipeline’s ability to retain k-mer files between runs facilitates rapid parameter adjustments without repeating earlier computations, as the k-mer counting is the most time consuming step. Our data suggest that a background-to-foreground ratio up to approximately five balances specificity and coverage effectively, as exemplified by the Sanger primer set achieving over a 95% specificity at a ratio of 5.81. When initial primer sets do not meet coverage goals, the force_target_threshold option can iteratively refine sets, though this process increases computational demands. With these steps, COATswga offers a robust platform for improving the detection and genotyping of *Plasmodium* and other pathogens.

## Conclusion

COATswga represents a substantial advancement in selective whole genome amplification (sWGA), providing an efficient and cost-effective approach for amplifying microbial genomes from complex field samples. Its enhanced capabilities in primer design, amplification uniformity, and sequencing accuracy, as demonstrated in *P. falciparum*, establish COATswga as a valuable tool for malaria genomics and microbial research more broadly. The new primer set for *P. falciparum* positions sWGA as an effective tool for improved malaria surveillance, drug resistance monitoring, and elimination efforts. As sWGA methodologies continue to advance, we anticipate further improvements to COATswga, such as refined primer selection algorithms and greater parallelization, that will expand its applicability. Past successes with sWGA across a range of microbes ensure that COATswga’s flexibility extends beyond *Plasmodium falciparum*, enabling enrichment of other pathogens, including bacterial and viral genomes, as well as other *Plasmodium* species. Currently, we are actively designing and testing COATswga primer sets for *Plasmodium vivax* and *Clostridium difficile*, aiming to broaden the software’s utility in infectious disease genomics.

To facilitate further optimization and applications of the sWGA method and related computational tools, the source code for COATswga can be found at https://github.com/bailey-lab/COATswga.

## Availability and Requirements

*Project name:* COATswga

*Project home page*: https://github.com/bailey-lab/COATswga

*Operating systems*: Platform independent

*Programming language*: Python

*Other requirements*: Conda environment, see project home page

*License*: GNU GPL

*Any restrictions to use by non-academics*: None

## Ethics approval and consent to participate

Not applicable

## Consent for publication

Not applicable

## Availability of data and materials

The source code and documentation for COATswga can be found at https://github.com/bailey-lab/COATswga. All data produced in the present study are available upon reasonable request to the authors.

## Authors’ contributions

KZ wrote the COATswga software, with advice from AS, AAF and JAB. KZ, AL, RD performed sWGA amplification, qPCR analysis, and Oxford Nanopore sequencing. AAF, AL and RD performed MIP sequencing. KZ and AAF executed data analysis and visualization. KZ and AAF wrote the manuscript, with contributions and edits from JAB. All authors read and approved the final manuscript.

## Acknowledgements

Not applicable

## Supplementary materials

### Supplementary Figures

**Figure S1:**
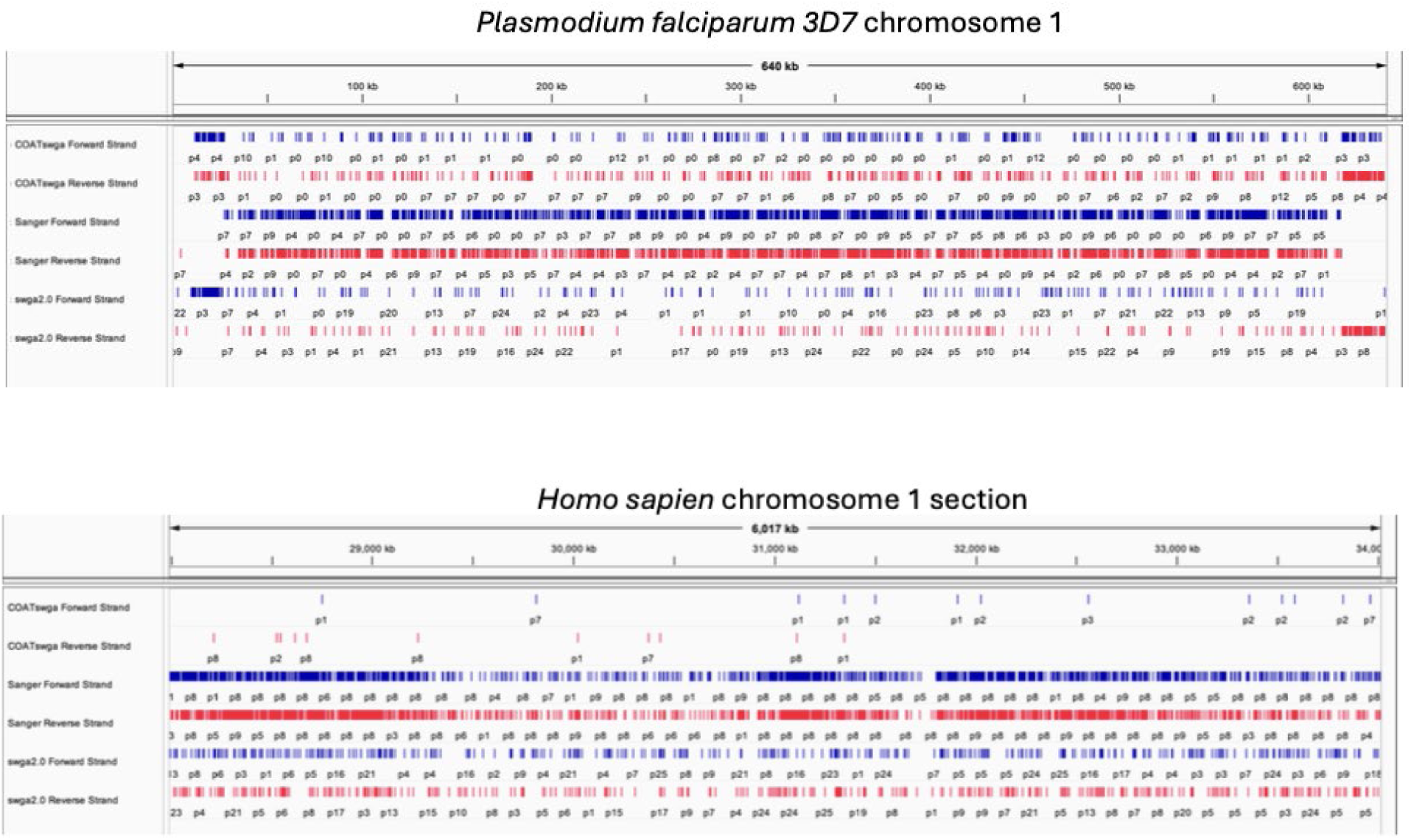
Representative screenshots of the Integrated Genomics Viewer (IGV) view of the hits for each primer set tested for the first chromosome of both the human and *Plasmodium falciparum* genomes. Blue and red indicate string matches in the forward and reverse strands of the genomes, respectively. Each tick indicates one hit at that point in the genome. In the order in the images, the tracks are: COATswga forward, COATswga reverse, Sanger set forward, Sanger set reverse, swga2.0 forward, swga2.0 reverse.

**Figure S2:**
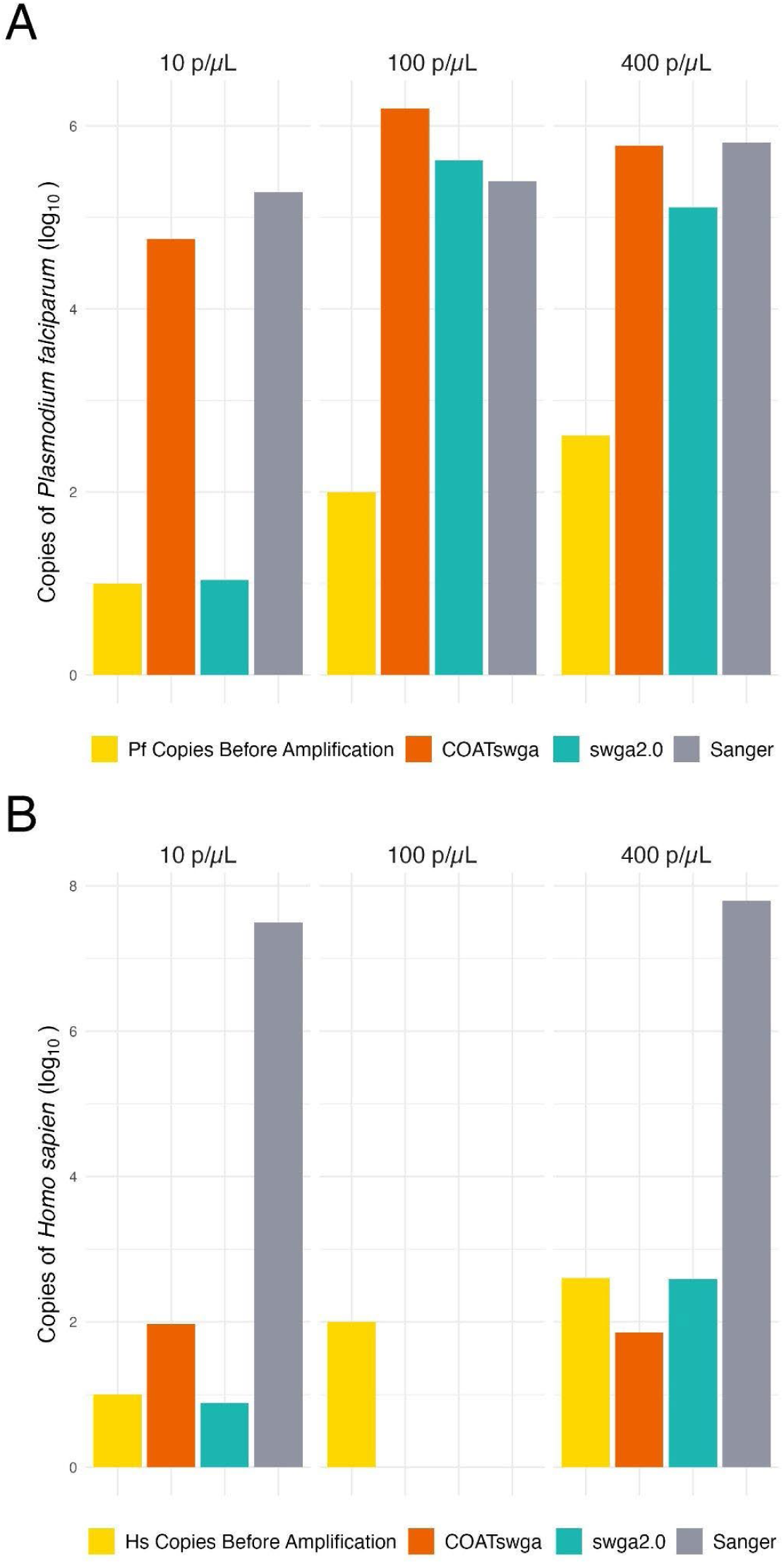
Bar plots of the qPCR amplification data of the three primer sets across the three levels of parasitemia in the (A) *Plasmodium falciparum* and (B) human genomes

**Figure S3:**
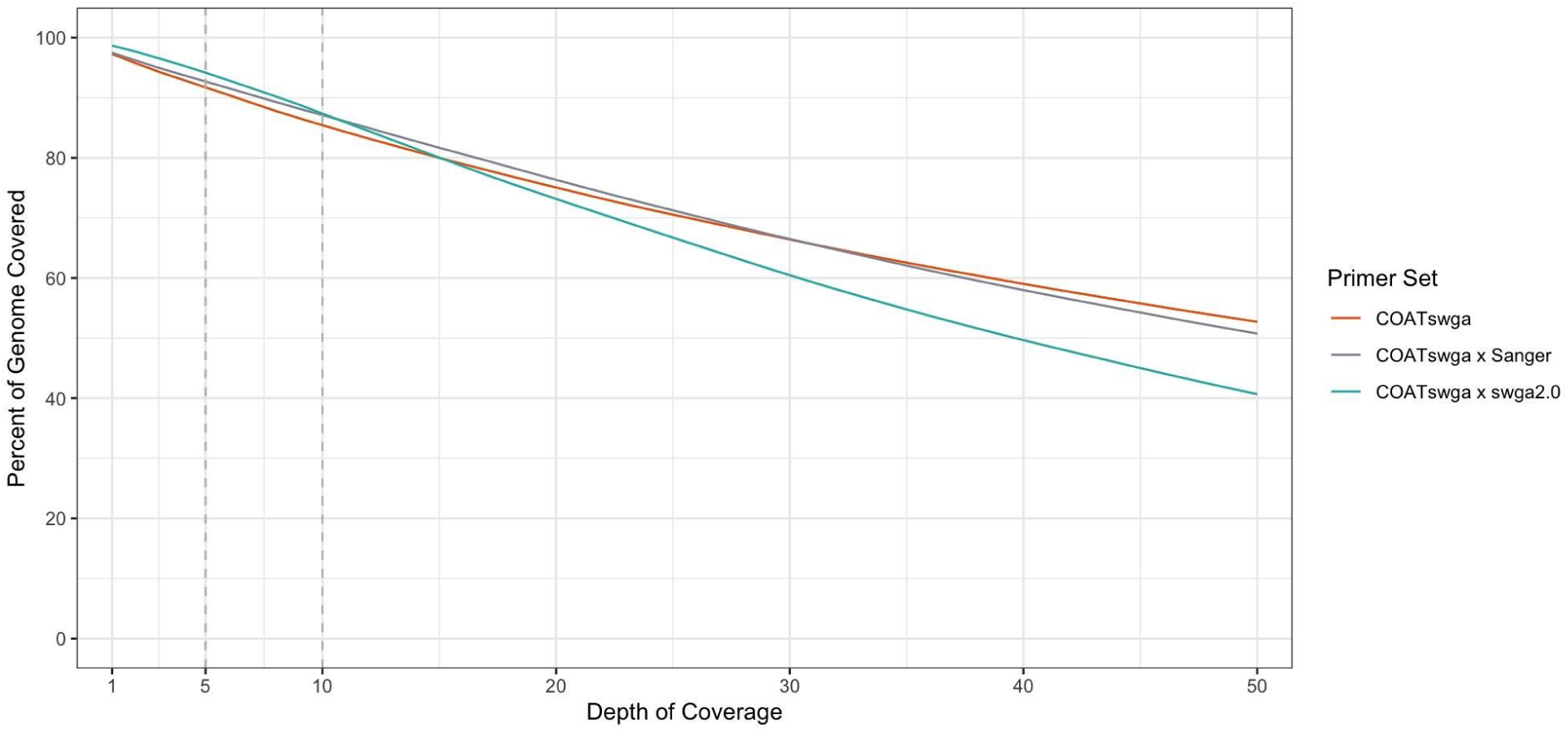
Coverage of whole *Plasmodium falciparum* genome with sequencing data from each primer pool combined bioinformatically. Each sample contains 2Gb of sequencing data. (COATswga = 2Gb, COATswga x Sanger = 1Gb COATswga + 1Gb Sanger, COATswga x swga2.0 = 1Gb COATswga + 1Gb swga2.0)

**Figure S4:**
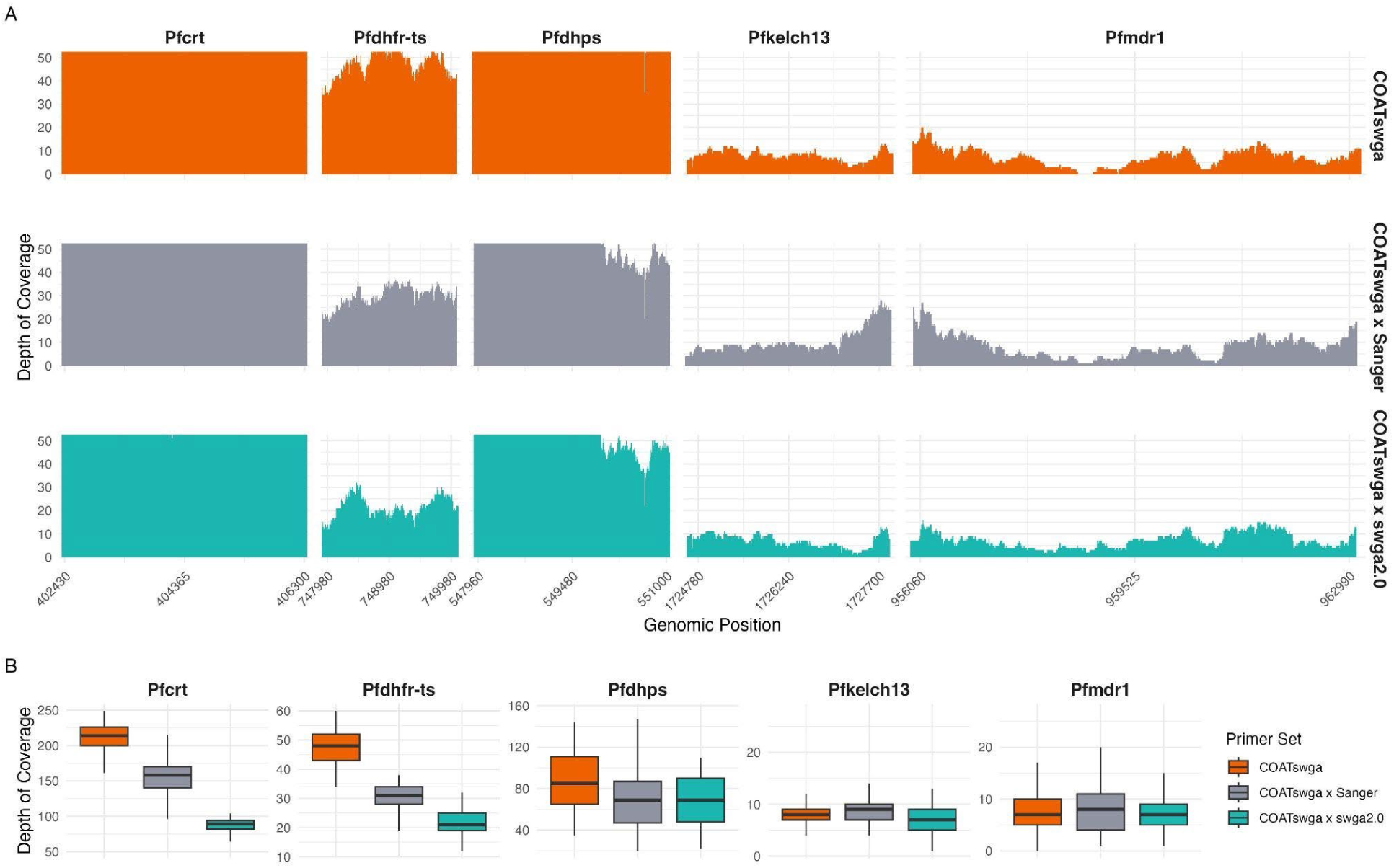
Coverage of drug resistance genes with sequencing data from each primer pool combined bioinformatically. Each sample contains 2Gb of sequencing data as in Figure S3.

**Figure S5:**
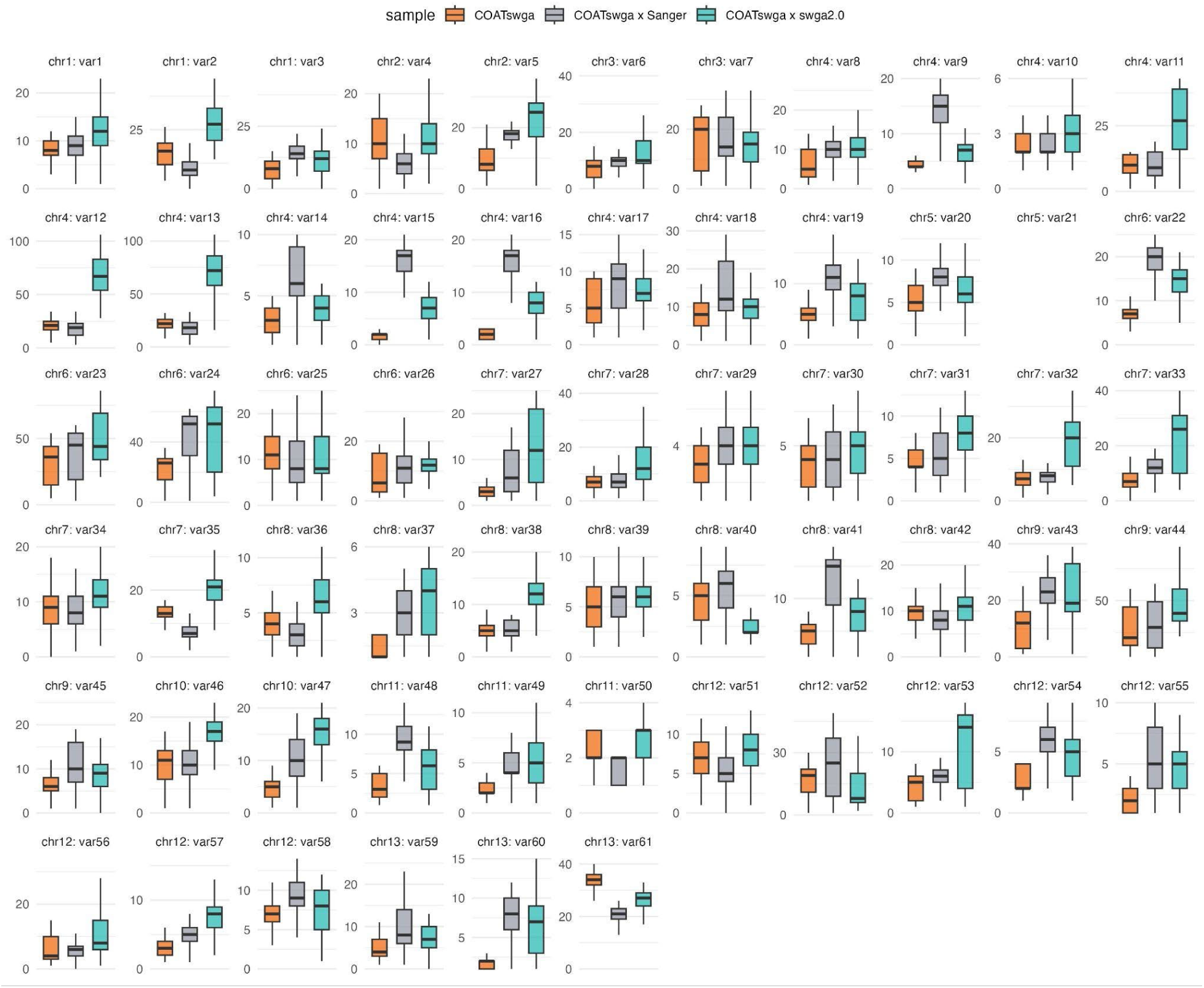
Coverage of var genes with sequencing data from each primer pool combined bioinformatically. Each sample contains 2Gb of sequencing data as in Figure S3 (COATswga = 2Gb COATswga, COATswga x Sanger = 1Gb COATswga + 1Gb Sanger, COATswga x swga2.0 = 1Gb COATswga + 1Gb swga2.0)

**Figure S6:**
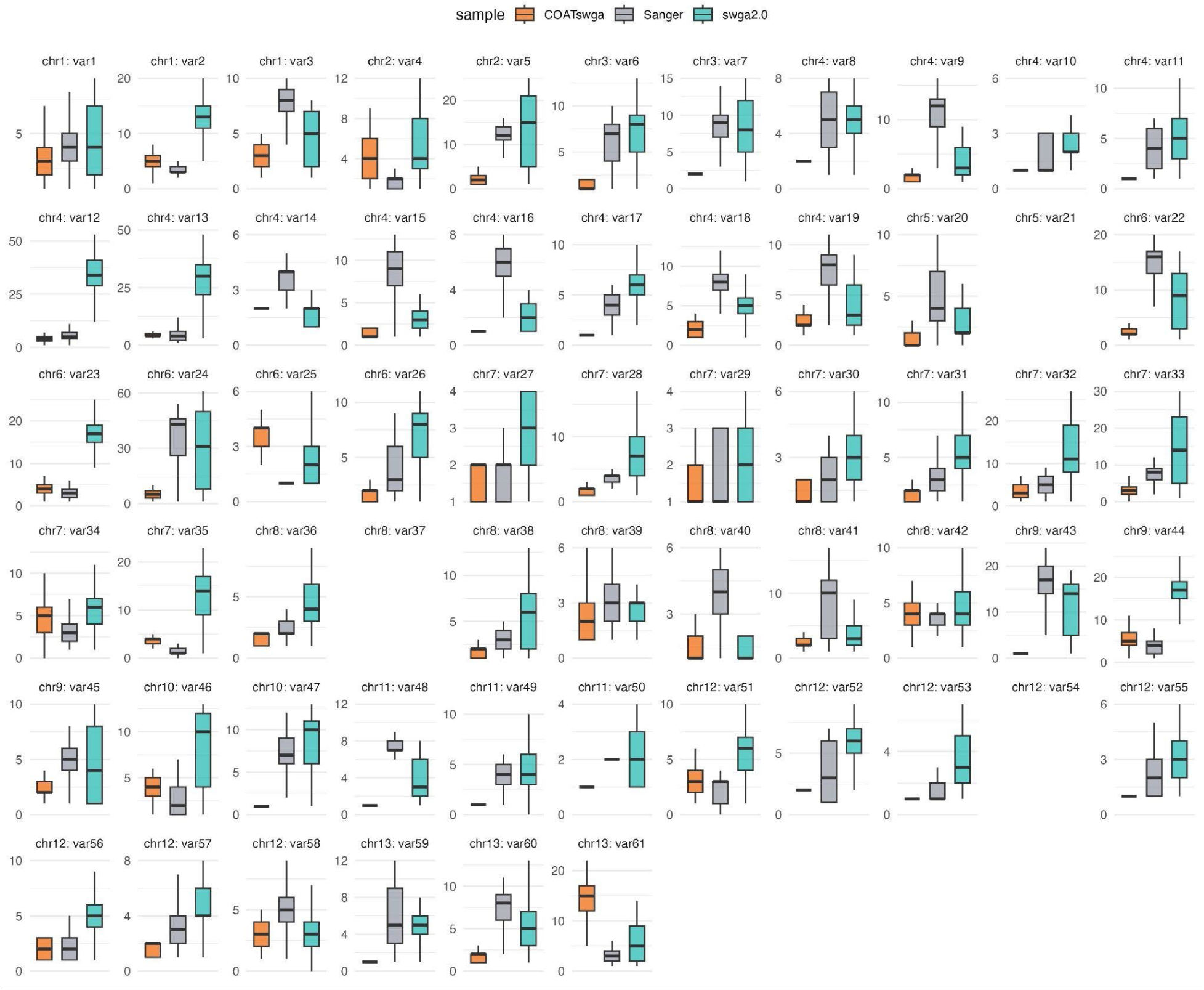
*var* gene coverage of each primer pool at 100 p/µL parasitemia

### Supplementary Tables

**Table S1:**
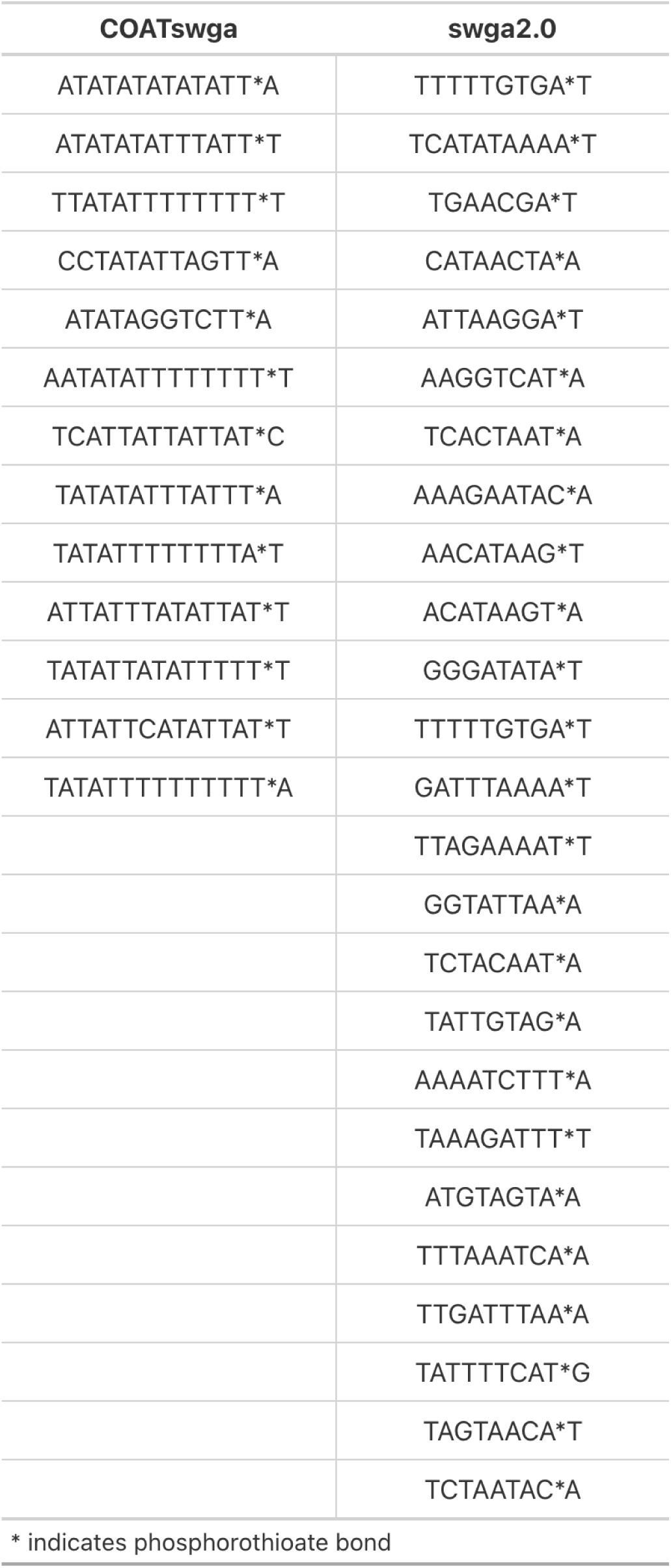
Primers contained in the COATswga and swga2.0 primer sets developed for this study.

**Table S2:** List of the 61*var* genes we used for our analysis and their locations in the *P. falciparum* genome.

| Chromosome | Start | End | Name | Strand |
| --- | --- | --- | --- | --- |
| chr1 | 29,510 | 37,127 | var1 | + |
| chr1 | 42,367 | 46,508 | var2 | - |
| chr1 | 607,390 | 614,894 | var3 | - |
| chr2 | 25,232 | 31,169 | var4 | + |
| chr2 | 916,352 | 923,649 | var5 | - |
| chr3 | 36,965 | 44,483 | var6 | + |
| chr3 | 1,030,822 | 1,038,255 | var7 | - |
| chr4 | 28,706 | 37,678 | var8 | + |
| chr4 | 45,555 | 56,861 | var9 | - |
| chr4 | 545,987 | 553,811 | var10 | - |
| chr4 | 561,667 | 569,343 | var11 | - |
| chr4 | 576,810 | 584,669 | var12 | - |
| chr4 | 591,949 | 599,850 | var13 | - |
| chr4 | 935,031 | 941,876 | var14 | - |
| chr4 | 946,169 | 953,774 | var15 | - |
| chr4 | 958,067 | 965,612 | var16 | - |
| chr4 | 969,031 | 976,592 | var17 | - |
| chr4 | 1,156,423 | 1,167,822 | var18 | + |
| chr4 | 1,173,053 | 1,180,227 | var19 | - |
| chr5 | 20,929 | 28,457 | var20 | + |
| chr5 | 1,333,470 | 1,342,965 | var21 | + |
| chr6 | 3,503 | 12,836 | var22 | + |
| chr6 | 18,586 | 22,722 | var23 | - |
| chr6 | 723,117 | 731,491 | var24 | - |
| chr6 | 1,353,946 | 1,366,431 | var25 | - |
| chr6 | 1,374,797 | 1,382,628 | var26 | - |
| chr7 | 20,307 | 28,084 | var27 | + |
| chr7 | 511,950 | 519,426 | var28 | - |
| chr7 | 527,338 | 535,132 | var29 | - |
| chr7 | 542,906 | 550,521 | var30 | - |
| chr7 | 552,158 | 559,079 | var31 | - |
| chr7 | 566,726 | 574,309 | var32 | - |
| chr7 | 581,386 | 588,924 | var33 | - |
| chr7 | 590,326 | 597,734 | var34 | - |
| chr7 | 1,417,588 | 1,426,235 | var35 | - |
| chr8 | 21,361 | 28,654 | var36 | + |
| chr8 | 29,700 | 39,034 | var37 | - |
| chr8 | 40,948 | 50,940 | var38 | + |
| chr8 | 431,165 | 439,052 | var39 | + |
| chr8 | 440,408 | 448,063 | var40 | + |
| chr8 | 459,312 | 466,568 | var41 | + |
| chr8 | 1,435,794 | 1,443,450 | var42 | - |
| chr9 | 20,080 | 27,886 | var43 | + |
| chr9 | 1,486,070 | 1,490,188 | var44 | + |
| chr9 | 1,495,579 | 1,503,337 | var45 | - |
| chr10 | 28,490 | 36,165 | var46 | + |
| chr10 | 1,642,401 | 1,649,949 | var47 | - |
| chr11 | 24,160 | 31,599 | var48 | + |
| chr11 | 32,666 | 42,387 | var49 | - |
| chr11 | 2,025,817 | 2,035,887 | var50 | + |
| chr12 | 16,973 | 24,498 | var51 | + |
| chr12 | 32,703 | 41,941 | var52 | + |
| chr12 | 46,788 | 56,806 | var53 | - |
| chr12 | 766,654 | 774,198 | var54 | - |
| chr12 | 1,694,150 | 1,703,088 | var55 | + |
| chr12 | 1,704,512 | 1,712,491 | var56 | + |
| chr12 | 1,719,574 | 1,727,457 | var57 | + |
| chr12 | 2,241,271 | 2,248,963 | var58 | - |
| chr13 | 21,364 | 28,788 | var59 | + |
| chr13 | 33,959 | 44,743 | var60 | - |
| chr13 | 2,884,786 | 2,892,341 | var61 | - |

**Table S3:**
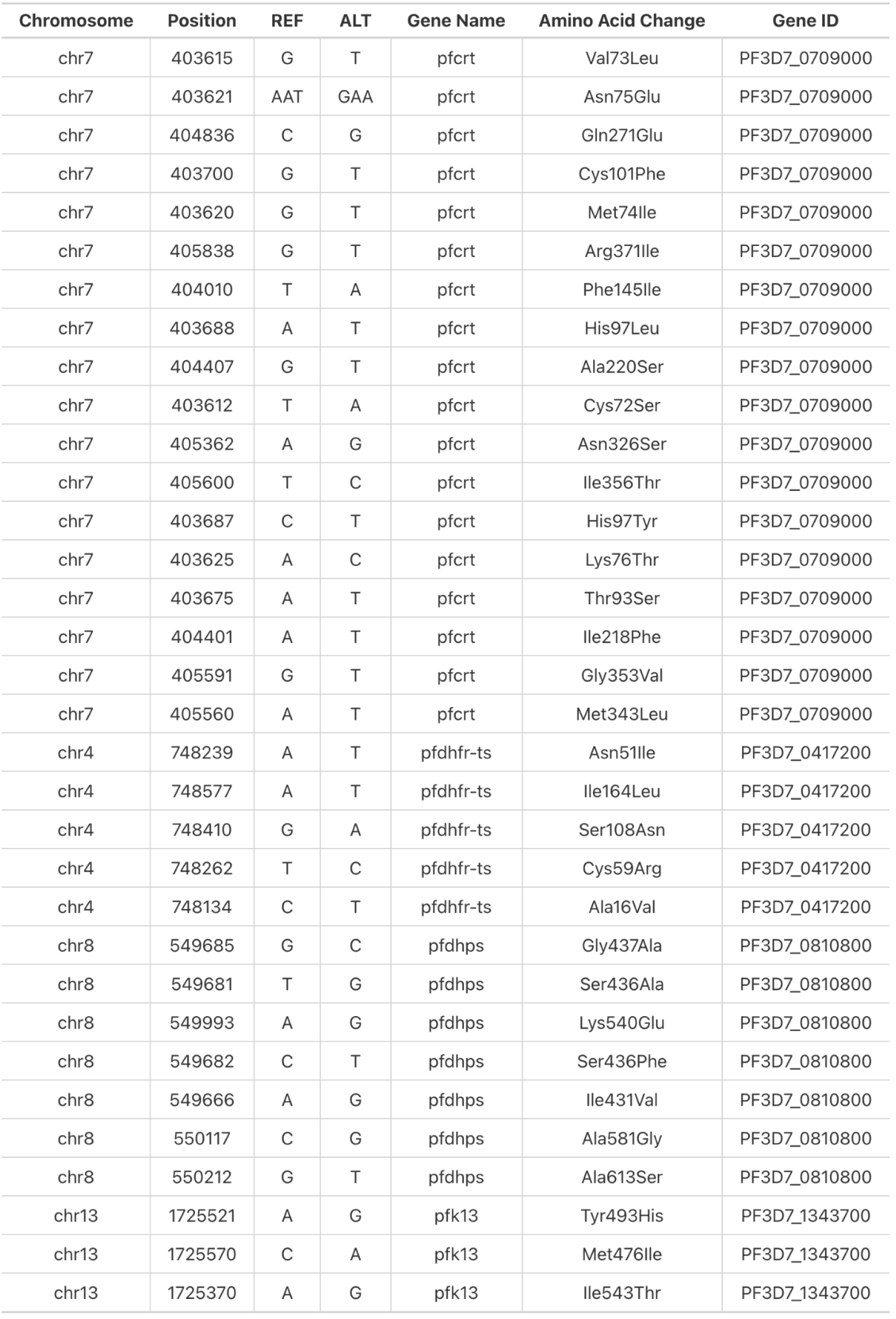
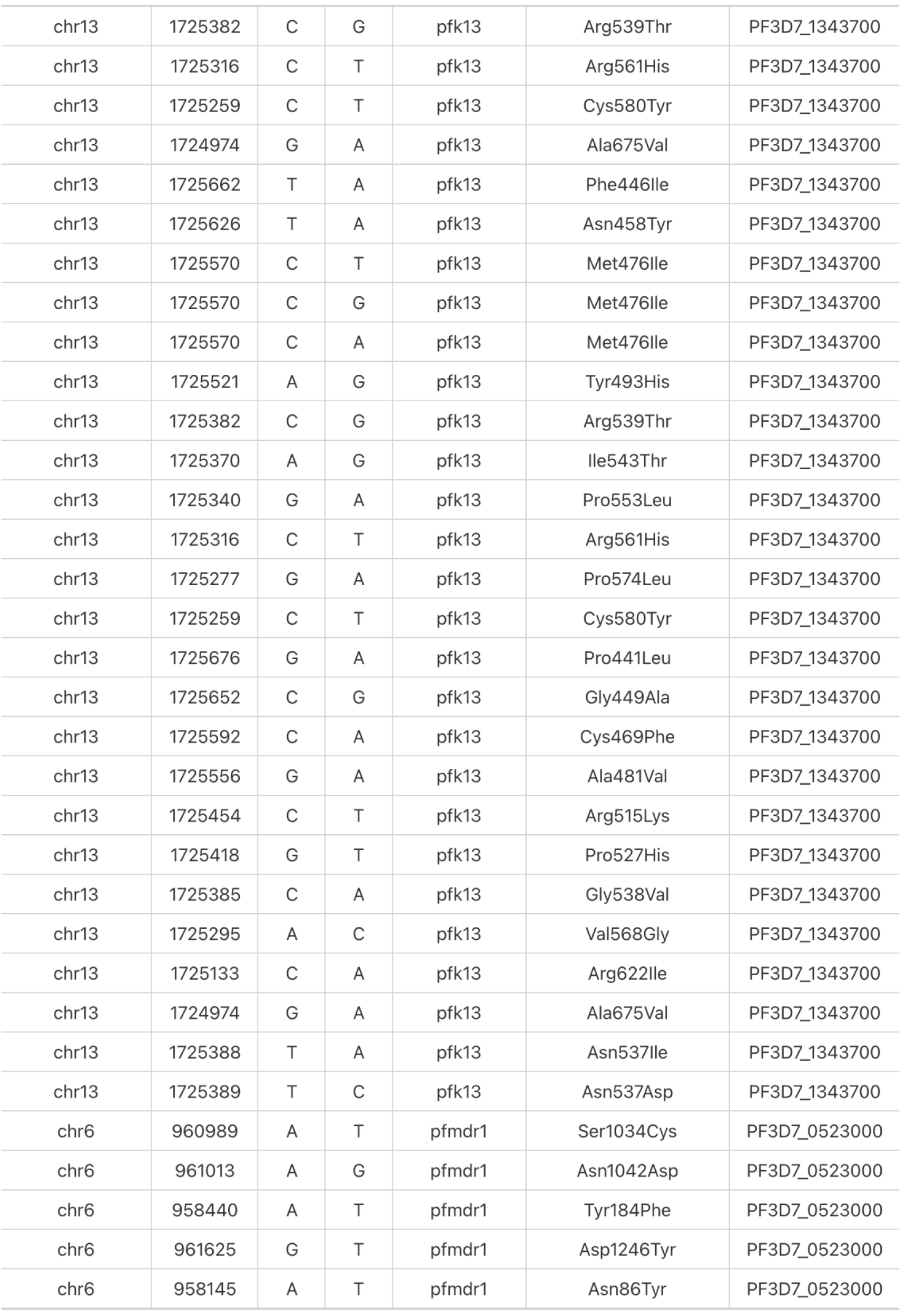
Key drug resistance loci covered in the *P. falciparum* genome in our COATsWGA testing with MIP and WGS sequencing.

